# Harnessing Stress: balancing the Burden of Slightly-Deleterious Variants by a Handicap

**DOI:** 10.1101/2024.01.25.577025

**Authors:** Victor Shamanskiy, Konstantin Popadin

## Abstract

Numerous empirical studies have revealed epistatic interactions among deleterious variants. In this paper, we assume such interactions are widespread and analyze the resulting shift in mutational burden and average population fitness following the introduction of a strong and universal stress (hereafter “handicap”). We demonstrate that organisms with a low burden of slightly-deleterious variants (SDVs) are more likely to survive exposure to a handicap, whether genetic or environmental, leading to a purifying effect on the population. We further discuss the potential applications of harnessing such interactions for evolutionary and population studies as well as for population management.

## Introduction

All organisms harbor numerous slightly-deleterious variants (SDV) in their genomes. However, the composition of this burden, as well as its total effect on fitness, remains poorly understood due to the system’s extreme complexity: thousands of SDVs, each with very small individual phenotypic effects, can interact with each other in non-additive ways, making the reconstruction of fitness from genomic data nearly impossible. For example, every human genome has approximately one thousand point SD coding variants, several thousand point SD regulatory variants, and dozens of SD copy number variants. Even though most of these mutations individually have small effects, their cumulative effect (mutation load) could substantially reduce the fitness of an individual. Despite its fundamental importance, the composition of this mutation load, as well as the total effect of the load on fitness, are poorly known due to extreme complexity of the system: thousands of SD mutations with very small individual phenotypic effects can interact with each other in non-additive ways. This makes the standard “bottom-up” approach (investigation of fitness consequences of each SD mutation in order to reconstruct the total mutation load) highly inefficient and laborious and even impossible if the fitness effects of SD mutations are too small to be measurable.

Considering the widespread negative epistatic interactions between SDVs (Sohail et al. 2017), we have formulated a “top-down” approach. This posits that a fit carrier of a severely-deleterious variant (hereafter ‘handicap’) is likely to have a reduced genome-wide burden of SDVs compared to controls (organisms without severely-deleterious variants). Recently, viewing trisomy 21 as a genetic handicap in humans (Popadin et al. 2018), we obtained empirical evidence supporting our hypothesis: live-born individuals with Down Syndrome exhibit a reduced burden of SDVs compared to control individuals. Here, we explore the handicap approach on a theoretical level: (i) Under what types of selection (e.g., hard/soft; varying strength and direction of epistasis) does the handicap approach work? (ii) How can we design experiments using populations of model organisms to estimate their mutation burden? (iii) It it possible to purify genomes from the burden of SDVs artificially? Answering these questions will aid in applying this approach to evolutionary and population studies to elucidate the mutational burden.

## Results

### 1. Handicap concept

If the biological fitness of an organism is determined as a function of its mutational load, and this mutational load consists mainly of unconditionally deleterious mutations, then we expect to observe a trade-off between the presence of a severely deleterious mutation and the total number of SDVs. We adopt the term ‘handicap’, introduced by Amotz Zahavi as a marker of high genome quality of a carrier (Zahavi 1975), and defined in the Oxford dictionary as “a circumstance that makes progress or success difficult.” We apply a “handicap principle”, where an organism bearing a severely-deleterious mutation (a ‘genetic handicap’) or one that has survived environmental stress (environmental handicap) is viable if its genome-wide load of SDVs is sufficiently low (Fig 1). The rationale for this hypothesis is that only highly fit organisms, with a low enough SDV load, can tolerate the effects of a handicap and survive.

**Fig 1.**
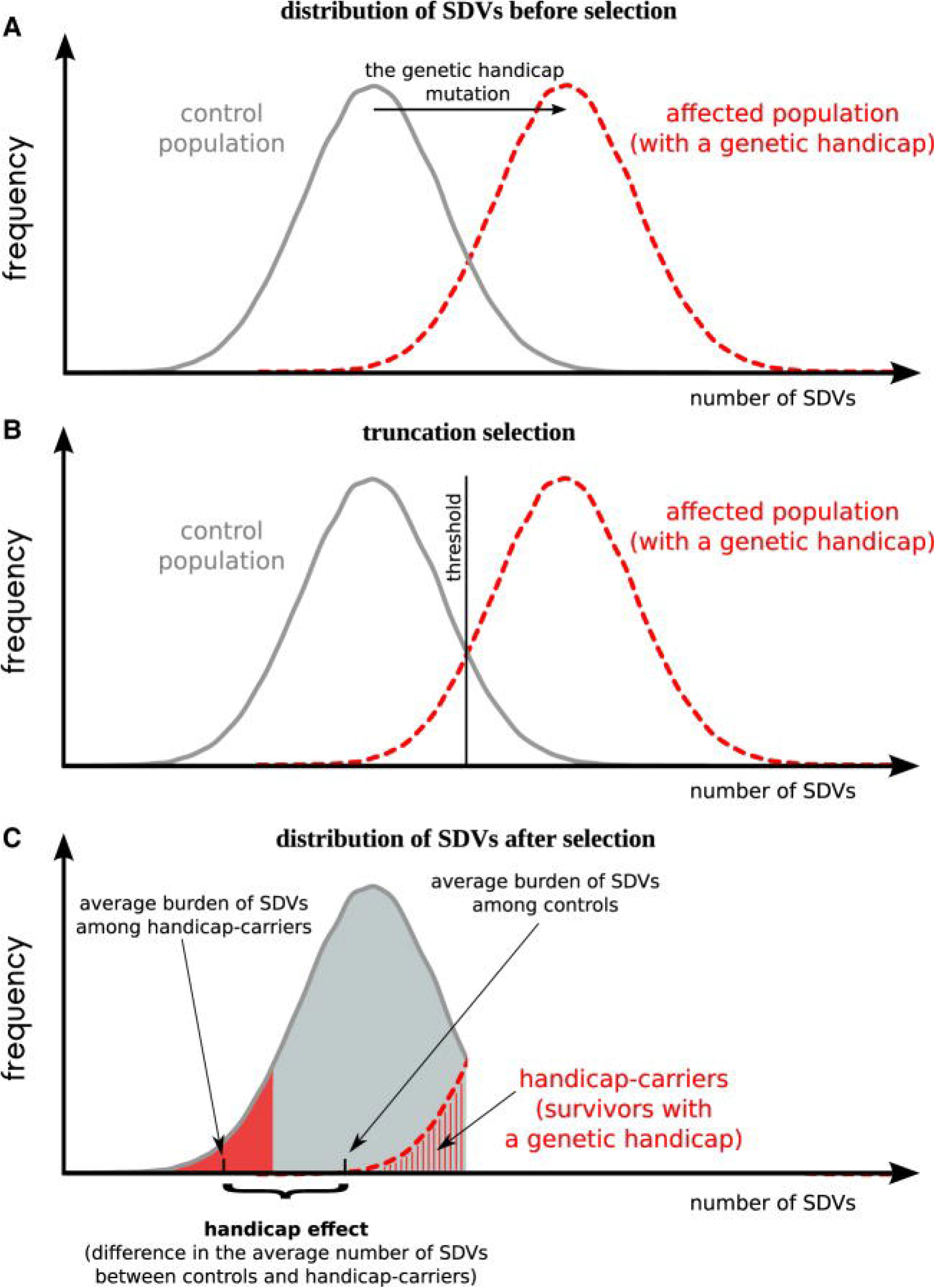
The handicap concept. (*A*) The distribution of the number of SDVs in control (gray) and affected (red) populations. The genetic handicap mutation (black arrow) is an equivalent of many SDVs. (*B*) Truncation selection eliminates all organisms with the number of SDVs higher than the given threshold (vertical black line) from both control and affected populations. (*C*) Handicap carriers have a decreased number of SDVs (SDVs do not include the genetic handicap per se) than controls; this difference represents the handicap effect. (Slightly deleterious genomic variants and transcriptome perturbations in Down syndrome embryonic selection, Popadin et al. 2018)

Here, we define a handicap as a severely-deleterious stress with a broad effect on cellular metabolism. The broad effect implies that the handicap is unlikely to be compensated or modified by a few conditional variants and therefore, might only be compensated by a genome-wide decreased load of unconditionally deleterious variants. In this scenario, we can approximate the total mutational load using the load of unconditionally deleterious mutations.

We assume here that the handicap is strong enough to divide the affected population into two groups: survivors and non-survivors, thus providing a straightforward classification of affected organisms based on their SDV loads (Fig 1). The main prediction of this hypothesis is that survivors will have a lower number of SDVs compared to controls (organisms survived without a handicap). Furthermore, the stronger the handicap’s severity, the greater the expected difference in SDV loads between survivors and controls.

The handicap approach offers two fundamental advantages. Firstly, the handicap, by causing negative epistasis with a load of SD variants, induces a very strong selection pressure that magnifies the fitness differences between organisms with high and low SDV loads – a potentially fundamental principle, which has been only poorly empirically demonstrated for any organism until now. Secondly, the severity of the handicap is expected to correlate with the difference in SDV loads between survivors and controls, allowing us to attribute a ‘handicap fitness effect’ to the quantity of SDVs in the controls but not in the survivors.

To validate this approach and to explore the interaction between a handicap and the load of SDVs, we pursue this question through mathematical and computational experiments.

### 2. Handicap - quantitative description

To quantitatively describe the effect of the handicap on the number of SDVs per individual, we use a classic model of selection against deleterious mutations (Charlesworth 1990)□. Assuming that all mutations have the same selective disadvantage, the difference in the mean number of SDVs before and after selection is given by:

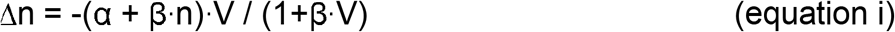

where α is the selection coefficient of deleterious mutations; β measures the strength of epistasis between deleterious mutations, n and V are the mean and variance of the number of SDVs before selection, respectively (Charlesworth 1990; Appendix 1)□.

Assuming that the handicap has an effect equivalent to that of k SDVs (handicap severity = k), the mean number of SDVs in a population is increased, but variance is unaffected. Thus, the change in the number of SDVs per individuals carrying a handicap follows as:

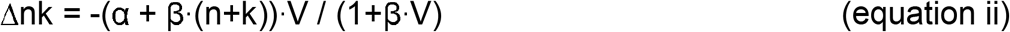

Hereafter, to approximate the handicap effect (H) as a difference in the efficiency of purifying selection with and without handicap, we subtract Δnk from Δn:

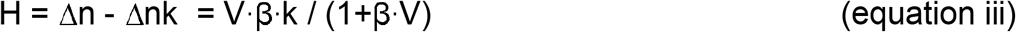

We see from equation (iii) that if there is no variance in the number of SDVs before selection (V=0), or if there is no epistasis between SDVs (β = 0), the handicap is absent (H = 0). However, when both the variance and the epistatic coefficient are higher than zero, the handicap effect increases linearly with handicap severity, k. Interestingly, when β approaches 1, indicating very strong positive epistasis, and variance increases infinitely, the handicap effect approaches the handicap severity:

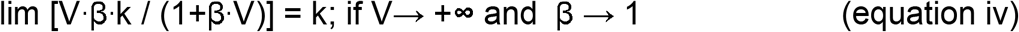

The interpretation of this result is that, under truncating selection and very high intra-population variation, individuals with handicap can survive only if they initially had n-k mutations, i.e. located in the left tail of the distribution (see Fig 1).

To derive a biologically meaningful range of handicap effects applicable to the human genome conditions, we produced two heatmaps. First (see Fig 2A), assuming that the average human genome harbors 1000 SDVs and the distribution is close to Poisson distribution (Sohail et al. 2017; Kondrashov 1995)□, we fixed the variance as 1000, fluctuated β from 0 to 0.1 (Charlesworth 1990)□, and applied a handicap severity value between 0 to 200, where 200 is the strongest difference in the SDVs between maximally and minimally loaded individuals from the population. This maximal handicap severity equals 200 assures us that there is a non-zero chance that an organism with a given handicap will be able to survive. We observed an increase in the handicap effect with both the handicap severity and the strength of epistasis. While it is intuitively expected that handicap severity is associated with the handicap effect, the strength of the negative epistasis is less obvious. To illustrate the importance of the negative epistasis we compared additive and negatively epistatic selection using simple example and revealed that only in epistatic scenario selection more effectively eliminates highly loaded genomes, thus significantly decreasing the mean number of SDVs among survivors.

**Fig 2.**
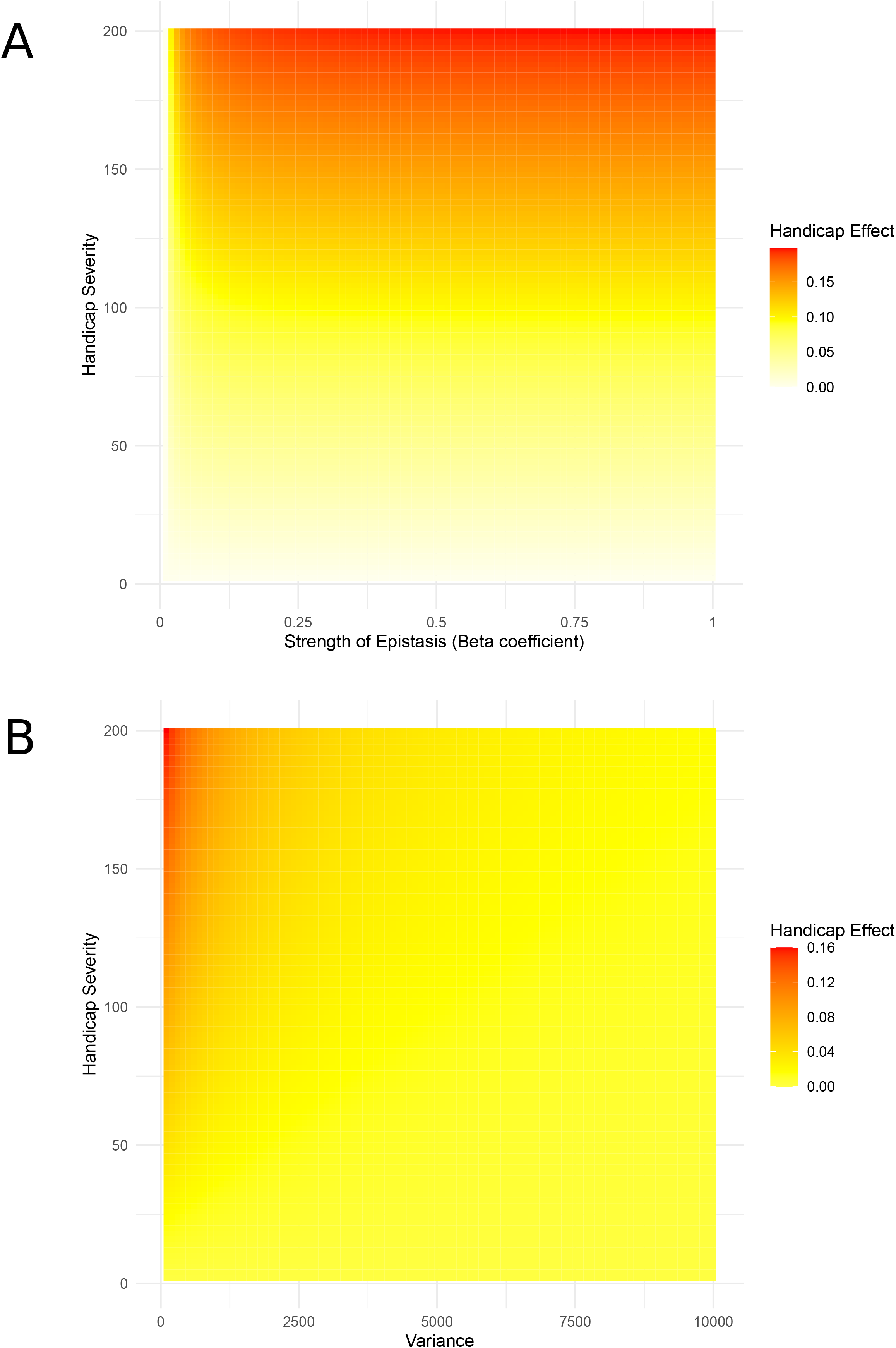
Heatmaps of handicap effect. **A**. With β fluctuated from 0 to 0.1 and the variance fixed as 1000. **B**. With β fixed as 0.0008 and the variance fluctuated from 0 to 10000

Second (see Fig 2B), we fixed β as 0.0008 (Charlesworth 1990)□ and fluctuated the variance from 0 to 10000 keeping the same range of handicap severity (0 to 200). In this scenario, the handicap effect increases with both the handicap severity and the variance of SDVs in a population.

Thus, the handicap effect, under biologically meaningful parameters, might be as strong as 20% of the total mutation load (Fig 2A – the strongest handicap effect is close to 200 out of 1000). Considering that negative epistasis is a predominant mode of interaction between SDVs (Sarkisyan et al. 2016; Sohail et al. 2017), especially in the case of organisms with complex genome (Sanjuán and Elena 2006)□ we conclude that the handicap principle is a universal concept applicable to model and non model species with complex genomes.

## Discussion

Here we propose that the negative fitness consequences of a severe event (genetic or environmental), which we call handicap, might be partially compensated by a reduced genome-wide load of SDVs. Thus, survivors after the exposition to a handicap are expected to have a decreased load of SDVs as compared to controls. Based on this concept it is possible to conduct an experiment to purify the genome of organisms exposing them to environmental handicap (https://doi.org/10.1101/2023.01.20.524958).

## Acknowledgments

VSh and KP were supported by Ministry of Science and Higher Education of the Russian Federation (agreement no. 075-15-2021-1084) for design of the experiment, performance of the experiment and analysis of the results.

## Notes

### Competing Interest Statement

The authors have declared no competing interest.

